# Individuals with epilepsy and a history of comorbid depressive symptoms exhibit disrupted fronto-temporo-limbic connectivity during cognitive conflict encoding

**DOI:** 10.1101/2024.10.10.617540

**Authors:** Aniruddha Shekara, Alexander Ross, Daniel J. Soper, Angelique C. Paulk, Sydney S. Cash, Alik S Widge, Paula K. Shear, John P. Sheehy, Ishita Basu

**Affiliations:** Department of Neurosurgery, University of Cincinnati College of Medicine, Cincinnati, OH 45267, USA; Department of Biomedical Engineering, University of Cincinnati College of Engineering and Applied Science, Cincinnati, OH 45219, USA; Department of Neurology, Center for Neurotechnology and Neurorecovery, Massachusetts General Hospital, Harvard Medical School, Boston, MA 02114, USA; Department of Psychiatry, University of Minnesota, Minnesota, MN 55455, USA; Department of Psychology, University of Cincinnati College of Medicine, Cincinnati, OH 45221, USA

**Keywords:** Executive dysfunction, functional connectivity, depression, stereotactic EEG, dynamic causal modeling, response conflict

## Abstract

**Background:** Depressive disorders are associated with impaired cognitive control, yet the neural mechanisms underlying these deficits remain poorly understood. We used stereotactic EEG (sEEG) in individuals with medically refractory epilepsy to determine whether a history of comorbid depressive symptoms was associated with altered behavioral and neural responses encoding cognitive conflict.

**Methods:** sEEGs were recorded from fronto-temporo-limbic regions in 28 participants (age: 35.4±8.9, female: 15/28) performing a Multi-Source Interference Task (MSIT). Psychiatric histories, neurobehavioral interviews, and symptom ratings were used to categorize participants with a history of depressive symptoms (EDS, n=13), or as epilepsy controls without depression history (n=15). Behavioral, spectral power, and coherence responses to conflict were determined by generalized linear mixed-effects models. Dynamic Causal Modeling (DCM) estimated theta–alpha (4-15Hz) effective connectivity between dorsolateral prefrontal cortex (dlPFC), dorsal anterior cingulate cortex (dACC), and lateral temporal lobe (LTL) during conflict.

**Results:** EDS participants demonstrated greater conflict-related slowing without difference in accuracy from controls. Conflict elicited distributed fronto-temporo-limbic oscillations in both groups. EDS participants showed greater conflict-related theta/alpha increases in PFC, LTL, and amygdala, and broadband increases in LTL power, and reduced left dlPFC-LTL theta/alpha coherence compared to controls. DCM model estimated effective connectivity from left dlPFC➔right LTL and left dACC➔left LTL was reduced in EDS.

**Conclusion:** Increased oscillatory responses in EDS may reflect a need for greater cognitive resources due to disrupted communication in fronto-temporo-limbic circuits encoding conflict. Our results highlight a role of these circuits in conflict encoding and can inform interventions for improving cognitive control in these populations.

## Introduction

Depressive disorders are a leading cause of disability and socioeconomic burden affecting approximately 8.5% of U.S. adults in the past year.^1^ Despite widespread availability, treatment outcomes for depression remain inconsistent, due in part to a lack of specificity for dysfunctional brain circuits.^2,3^ High comorbidity among depressive disorders reduces therapeutic efficacy,^4^ while symptom-based assessments may not fully capture individual heterogeneity. An alternative strategy would be to target functional deficits defined by the Research Domain Criteria (RDoC)^5^ that are associated with compromised neural circuits.

Cognitive control, which enables flexibly adapt thoughts and behaviors to achieve longitudinal goals, is frequently impaired in depressive disorders.^6,7^ Cognitive control coordinates attention, working memory, perception, and response inhibition to suppress immediate actions in favor of goal-directed ones.^8,9^ Deficits in response inhibition often manifest in depression as maladaptive, repetitive negative thoughts and behaviors.^10,11^ Thus, cognitive control can serve as a clinically relevant target for therapeutic intervention.

Response inhibition can be assessed by tasks such as the Stroop test,^12^ which introduce cognitive conflict between task-relevant and task-irrelevant information. Conflict evokes theta (4-8Hz) and high-gamma (70-110Hz) oscillations in prefrontal cortex (PFC) and dorsal anterior cingulate cortex (dACC), which play key roles in conflict detection and response inhibition.^13-16^ Recent stereotactic EEG (sEEG) studies have examined these oscillatory signatures with greater spatial resolution. Studies of sEEG activity during a multi-source interference task (MSIT) eliciting robust neural responses to conflict^17-19^ have observed distributed conflict encoding in the orbitofrontal cortex (OFC), lateral temporal lobe (LTL), amygdala, and hippocampus, in addition to PFC and dACC.^20-23^ While the LTL is integral to semantic memory, speech, emotion, and sensory processing,^24-28^ its role in cognitive control remains unclear. Epilepsy patients, who comprise the majority of intracranial cognitive control studies, frequently demonstrate LTL impairment and cognitive dysfunction,^29^ which suggests the LTL may play a significant role in cognitive control. Furthermore, patients with epilepsy frequently experience depression,^30,31^ warranting investigation into how comorbid depressive symptoms alter cognitive control in this population.

Extant literature implicates cognitive control deficits in depression arise from aberrant PFC and dACC responses during conflict, rather than overt behavioral impairment. Some studies indicate heightened PFC and dACC activity reflect inefficient cognitive processing,^32-35^ while others report PFC/dACC hypoactivity associated with impaired conflict resolution.^36^ Together, these findings suggest depression may lead to deleterious and/or compensatory changes in cognitive control circuitry, though the extent of these deficits remains unclear. Importantly, associations between conflict encoding and depressive symptoms have yet to be examined with sEEG, which can capture large-scale neural interactions with high spatiotemporal resolution.

Previously, we reported that conflict enhances prefrontal theta power on sEEG during the MSIT,^22^ but did not consider neuropsychiatric differences in oscillatory responses. Here, we examined fronto-temporo-limbic oscillations encoding conflict in epilepsy participants with history of comorbid depressive symptoms (EDS), and a non-depressed epilepsy control group. Our goal was to determine whether EDS participants exhibit altered spectral responses and function/effective connectivity in oscillatory networks engaged during cognitive control (Fig. 1A). Given that depressed individuals often perform comparably to controls on cognitive control tasks,^37^ we hypothesized that EDS participants would demonstrate altered spectral responses and connectivity within fronto-temporo-limbic networks encoding conflict while maintaining comparable task performance to controls.

**Figure 1.**
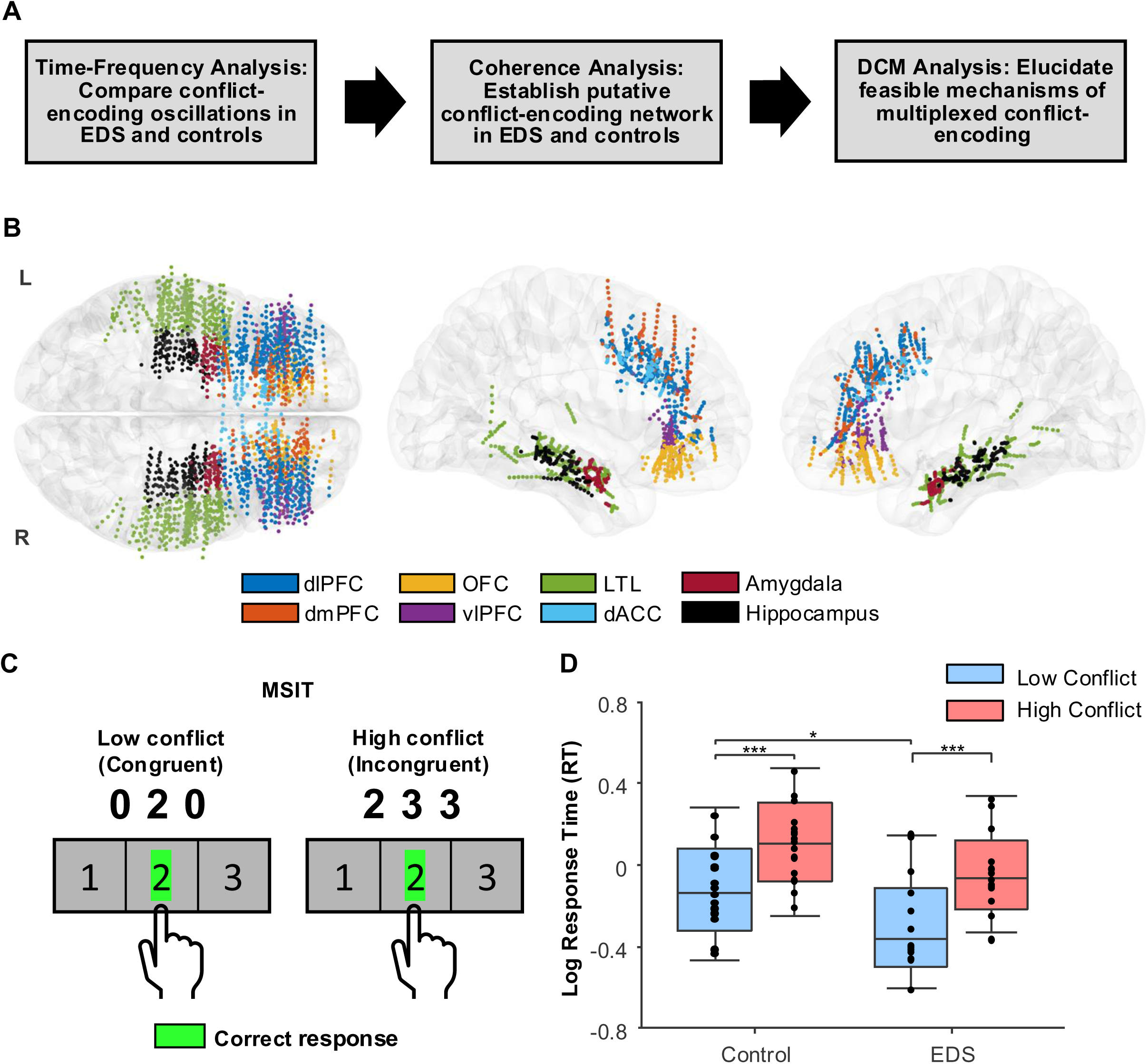
Experimental paradigm and response times (RT) during the MSIT. **(A)** Flowchart of analyses performed to characterize conflict-encoding networks in control and EDS participants. **(B)** sEEG electrode placements of all participants plotted on a standard brain template. **(C)** Schematic of the MSIT where participants must inhibit pre-potent responses on 50% of trials. **(D)** Log-transformed RT during low and high conflict trials. Central lines are median, bottom and top edges are 25% and 75%, and whiskers denote mean±SD. Markers represent average log-RT of individual participants. *p<0.05, ***p<0.001 (FDR-corrected).

## Materials and methods

### Participants

We analyzed sEEG data from 28 human participants with intractable epilepsy (age: 35.4±8.9, female: 15/28) implanted with stereotactic depth electrodes (Ad-Tech Medical or PMT) for seizure localization (Fig. 1B, Supplemental Methods). Seventeen participants (age: 34.4±10.4, female: 9/17) were previously recruited at Massachusetts General Hospital and Brigham and Women’s Hospital (MGH/BWH),^22^ and eleven participants (age: 36.9±6.0, female: 6/11) were recruited at the University of Cincinnati Medical Center (UCMC) during the study period. Study procedures took place in the epilepsy monitoring unit (EMU) at least two days post-implantation. Implantation procedures were performed as clinically indicated without research consideration. Written informed consent was obtained from study participants according to the Declaration of Helsinki, and study procedures were approved by MGH/BWH and UCMC Institutional Review Boards.

### Neuropsychiatric assessment and participant categorization

Psychiatric histories were obtained from neurobehavioral interviews with a clinical neuropsychologist prior to or during monitoring, and chart reviews (UCMC). Nineteen participants additionally completed the Beck Depression Inventory-II (BDI-II)^38^ or Patient Health Questionnaire-9 (PHQ-9)^39^ during their interview. Given the heterogeneity in clinical data across study sites, we classified participants as EDS if they met one more of the following consensus criteria: (1) assessed as having current or prior depressive symptoms by a neuropsychologist, (2) prior diagnosis of a unipolar depressive disorder, or (3) endorsed moderate or greater symptoms on the BDI-II (≥20) or PHQ-9 (≥10).

### Multi-Source Interference Task (MSIT)

Participants performed a version of the MSIT^17,40^ (Fig. 1C) at MGH/BWH as previously described,^22^ or on a laptop at UCMC (https://github.com/neuromotion/task-msit). MSIT stimuli consisted of three numbers between 0-3, one of which was unique. Participants were instructed to press the number key (1, 2, or 3) corresponding to the identity, but not position, of the unique number. During low conflict trials, the identity and position of the unique number aligned, and flankers were invalid (“0”). During high conflict trials, the identity of the unique number differed from its position on the keyboard, and flankers were valid responses. Stimuli appeared for up to 2s, followed by a fixation cross randomly jittered between 2-4s. Participants were instructed to respond as quickly and accurately as possible. Each participant completed a training block, then 2-8 blocks of 48 or 64 trials. Training data were excluded from analysis.

### sEEG acquisition and preprocessing

Neural data were recorded from sEEG electrodes placed in frontal, temporal, and subcortical regions. Recordings at MGH/BWH were acquired using a Neural Signal Processor (Blackrock Microsystems) sampling a 2kHz, and at UCMC using a Natus Quantum system (Natus Medical Inc.) sampling at 512Hz. Electrodes localized to dorsolateral prefrontal cortex (dlPFC), dorsomedial prefrontal cortex (dmPFC), ventrolateral prefrontal cortex (vlPFC), OFC, LTL, dACC, amygdala, and hippocampus were selected for analysis (Table S1). Data were bipolar re-referenced, notch-filtered, down-sampled to 512Hz if necessary, and inspected for artifacts (Supplemental Methods).

### Electrode localization

Electrode localization was performed using a modular FreeSurfer-based pipeline.^41,42^ Preoperative T1-weighted MRI scans were manually registered in FreeSurfer to postoperative CT scans of implanted electrodes mapped to the DKT40 atlas^43^ using an automated probabilistic labeling algorithm.^44^ For this study, we considered regions sampled in ≥5 participants within each group. Using these criteria, we selected electrodes localized to the following regions defined from atlas labels: left and right dorsolateral prefrontal cortex (dlPFC), dorsomedial prefrontal cortex (dmPFC), ventrolateral prefrontal cortex (vlPFC), OFC, LTL, dACC, amygdala, and hippocampus (Table S1). Electrodes outside of selected regions were excluded from analyses.

### Time-frequency analysis

Stimulus-locked time-frequency power was estimated within theta (4-8Hz), alpha (8-15Hz), beta (15-30Hz), gamma (30-55Hz), and high-gamma (70-110Hz) bands using Morlet wavelet decomposition. Single-trial power was averaged during conflict encoding (0.1s to trial response time) and normalized to baseline (-0.5 to -0.1s preceding stimulus onset) by frequency band using a log ratio. Spectral coherence^45^ was estimated within theta, alpha, and beta bands between electrode pairs, and averaged during conflict encoding (0.1s to median response time) and baseline for low and high conflict trials (Supplemental Methods). Coherence during conflict encoding were Fisher z-transformed, and normalized by the difference from baseline.^46^

### Dynamic Causal Modeling (DCM)

DCM is a Bayesian framework to make inferences on effective connectivity by parameterizing biophysical models of hidden neuronal states that predict observed data features.^47^ Model parameters are estimated using a Variational Bayes procedure that maximizes the log evidence for observing the data under a given model architecture. Additionally, DCM allows for testing hypotheses on how task conditions modulate connectivity parameters. Based on previous literature on conflict-encoding networks in healthy and depressed participants,^48-50^ results from spectral GLMEs, and regional sampling in our dataset, we selected the left dlPFC, left dACC, left LTL, and right LTL as regions within a putative conflict-encoding network for DCM analysis.

We used DCM for cross-spectral densities (CSD) as implemented in SPM12^51,52^ to model theta-alpha (4-15Hz) cross-spectra within this network. Regional activity was modeled by Jansen-Rit neural mass models of local field potentials (LFP)^53^ with extrinsic (inter-regional) and intrinsic (within-region) connectivity parameters. Forward connections from LTL➔dlPFC/dACC and backward connections from dlPFC/dACC➔LTL were defined from hierarchy in sensory/perceptive and cognitive processing, and evidence for prefrontal regulation of dACC/dmPFC during conflict.^49,54,55^ Reciprocal lateral connections were included between left and right LTL. We included modulatory effects of conflict on all specified extrinsic and intrinsic connections.

DCMs were estimated for a subset of 15 participants with electrode coverage in all selected regions. To reduce data dimensionality while retaining information on inter-regional variance, single value decomposition (SVD) was performed on electrode data for each participant and region.^56^ Single-trial data of principal eigenvariates obtained from SVD were used as representative signals for each region in the DCM analysis. We estimated two DCM models for each participant during baseline (-0.5 to -0.1s) and conflict encoding (+0.1s to median high conflict response time) task windows.

### Statistical analysis

We used generalized linear mixed-effects models (GLME) to estimate conflict (low, high) and group (control, EDS) effects on behavior and neural responses during MSIT performance. GLMEs modeled single-trial response data while accounting for inter-participant/inter-electrode variance, imbalanced sampling, and missing/artifactual trials. We used normal or log-normal distributions and an identity link function for all GLME analyses based on approximated data distributions (Supplemental Methods).

#### Behavior analyses

A Wilcoxon rank-sum test was used to compare accuracy between groups, after which all following analyses focused on trials with correct responses. Single-trial response times (RT) were log-transformed to approximate normality, then fit to a GLME (*fitglme)* of conflict and group coded as binary fixed-effects, and a random-effect of participant:

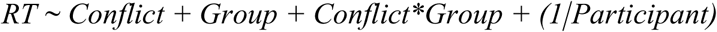

#### Spectral analyses

Recordings from 2981 bipolar-referenced channels in fronto-temporo-limbic regions during 5891 correct trials were retained for neural analyses following artifact rejection (278 trials, or 4.45%). GLMEs were fit to single-trial normalized power during conflict encoding for each region and frequency band (theta to high-gamma) with participant and channel nested random effects:

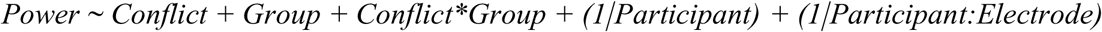

Significance of model predictors was determined using Wald tests on GLME coefficients. Significant interaction terms were interpreted post-hoc by Wald tests on coefficient contrasts (conflict: high>low, group: EDS>control). Models without significant interaction terms were reduced in subsequent GLMEs to estimate main effects of conflict and group. P-values were corrected for false discovery rate (FDR) by a step-up procedure (*fdr_bh^57^*) with q=0.05.

We modeled baseline-normalized coherence within theta, alpha, and beta bands between regions where conflict modulated band power in one or both groups (significant high>low conflict contrast or main effect). GLMEs were fit to normalized coherence during conflict encoding between electrode pairs within selected regions for each frequency band, with nested random effects of participant and electrode pair:

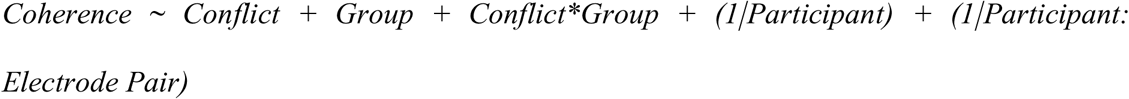

Post-hoc analyses and model reduction procedures were performed as for power models.

#### Parametric Empirical Bayes (PEB)

PEB was used to estimate group differences in DCM effective connectivity parameters while accounting for uncertainty in parameter estimates and model architecture between participants as random effects.^58,59^ Connectivity parameters from baseline and conflict-encoding DCMs were fit to a PEB model with binary fixed-effects of group (control, EDS), task (baseline, conflict encoding), and a group x task window interaction coding for group differences in baseline-corrected connectivity estimates (conflict encoding>baseline):

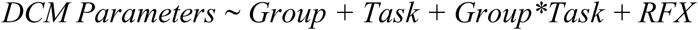

Post-hoc PEB models estimating baseline-corrected effective connectivity parameters were performed separately for control and EDS groups:

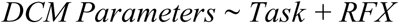

PEB models were optimized by an exhaustive greedy-search procedure to prune connectivity parameters that did not contribute to model evidence for the observed data and group/task effects.^59^

## Results

### Demographic data

MGH/BWH and UCMC participants did not significantly differ in age (t(28)=0719, p=0.479), sex (*χ*^2^=0.007, p=0.934), or handedness (Fisher exact test: p=0.085), and were pooled for all analyses (Table S2). Of the 28 participants, 13 met EDS criteria from psychiatric history, neurobehavioral interviews, and BDI-II/PHQ-9 scores (Table 1). Fifteen participants with no depressive symptom history and/or minimal/mild depression symptom ratings were designated as controls.

**Table 1.** Demographic and clinical characteristics of MGH/BWH and UCMC participants. Depression history and symptom severity varied across EDS participants and were often comorbid with anxiety or other psychiatric history. One participant with a history of premenstrual dysphoric disorder was designated to the control group based on a minimal BDI-II score. Control and EDS participants did not significantly differ in age (t(28)=0.457, p=0.651), sex (χ^2^=0.619, p=0.431), handedness (Fisher exact test: p=1), or site (χ^2^=1.152, p=0.283).

|  | **Control (n=15)** | **EDS (n=13)** |
| --- | --- | --- |
| Age (mean±SD) | 34.7±9.9 | 36.2±7.86 |
| Sex, female (n) | 7 | 8 |
| Handedness, right (n) | 11 | 11 |
| Site |  |  |
| MGH/BWH | 6 | 11 |
| UCMC | 7 | 4 |
| Psychiatric history (n) |  |  |
| Depressive disorder | 1 | 11 |
| Anxiety disorder | 1 | 11 |
| Other psychiatric disorder | 1 | 3 |
| Not available | 0 | 1 |
| BDI-II or PHQ-9 (n) |  |  |
| Minimum/Mild | 10 | 1 |
| Moderate/Severe | 0 | 7 |
| Not available | 5 | 5 |
| BAI or GAD-7 (n) |  |  |
| Minimum/Mild | 9 | 3 |
| Moderate/Severe | 0 | 5 |
| Not available | 6 | 5 |

### Behavioral data

We first compared MSIT accuracy and response times between groups. RT and accuracy were obtained for 6530 MSIT trials, after which 6169 correct trials (361/5.53% error or omitted) were retained for analysis. Mean accuracy on the MSIT was 94.49±6.47% and did not differ between EDS and controls (z=1.155, p=0.248). A GLME of conflict (low, high) and group (control, EDS) effects on log-normalized RT found a significant conflict x group interaction (β=0.032, p=0.002) (Table 2). Post-hoc contrasts showed that EDS participants demonstrated greater conflict-related slowing and faster responses on low conflict trials compared to controls (Fig. 1D).

**Table 2.** Effects of Conflict and Group on RT. Summary of post-hoc GLME coefficient contrasts estimating conflict (high>low) effects on RT in control and EDS participants, and group (EDS>control) differences in RT during low and high conflict trials. Bolded p-values denote effects that survived FDR-correction.

| **Contrast** | **$\beta$** | **F** | **p** |
| --- | --- | --- | --- |
| **Conflict (high>low)** |  |  |  |
| Control | 0.226 | 1128.911 | **<0.001** |
| EDS | 0.257 | 1176.610 | **<0.001** |
| **Group (EDS>control)** |  |  |  |
| Low Conflict | -0.173 | 5.647 | **0.017** |
| High Conflict | -0.141 | 3.770 | 0.052 |

### sEEG data

#### Fronto-temporo-limbic spectral responses encoding conflict in EDS and controls

We fit GLMEs to pre-response spectral power within theta to high-gamma bands in fronto-temporo-limbic regions to compare oscillatory responses encoding conflict between control and EDS. Significant conflict x group effects were observed across theta-gamma bands in left dlPFC, dACC, and hippocampus, right dmPFC, vlPFC, and amygdala, and bilateral LTL (Figs. 2A-B, Table S3). Post-hoc contrasts revealed greater increases in left LTL theta-beta power in EDS compared to controls during conflict (Fig. 2C, Table 3). EDS participants showed conflict-related increases in bilateral PFC, left dACC, and right amygdala theta/alpha power and broadband right LTL power, and reductions in gamma/high-gamma power in right dlPFC and LTL. In controls, beta power was increased in right vlPFC and reduced in right LTL by conflict. Group contrasts during high conflict found greater left hippocampus gamma power in EDS compared to controls (β=0.024, FDR-p=0.007) (Table S4).

**Figure 2.**
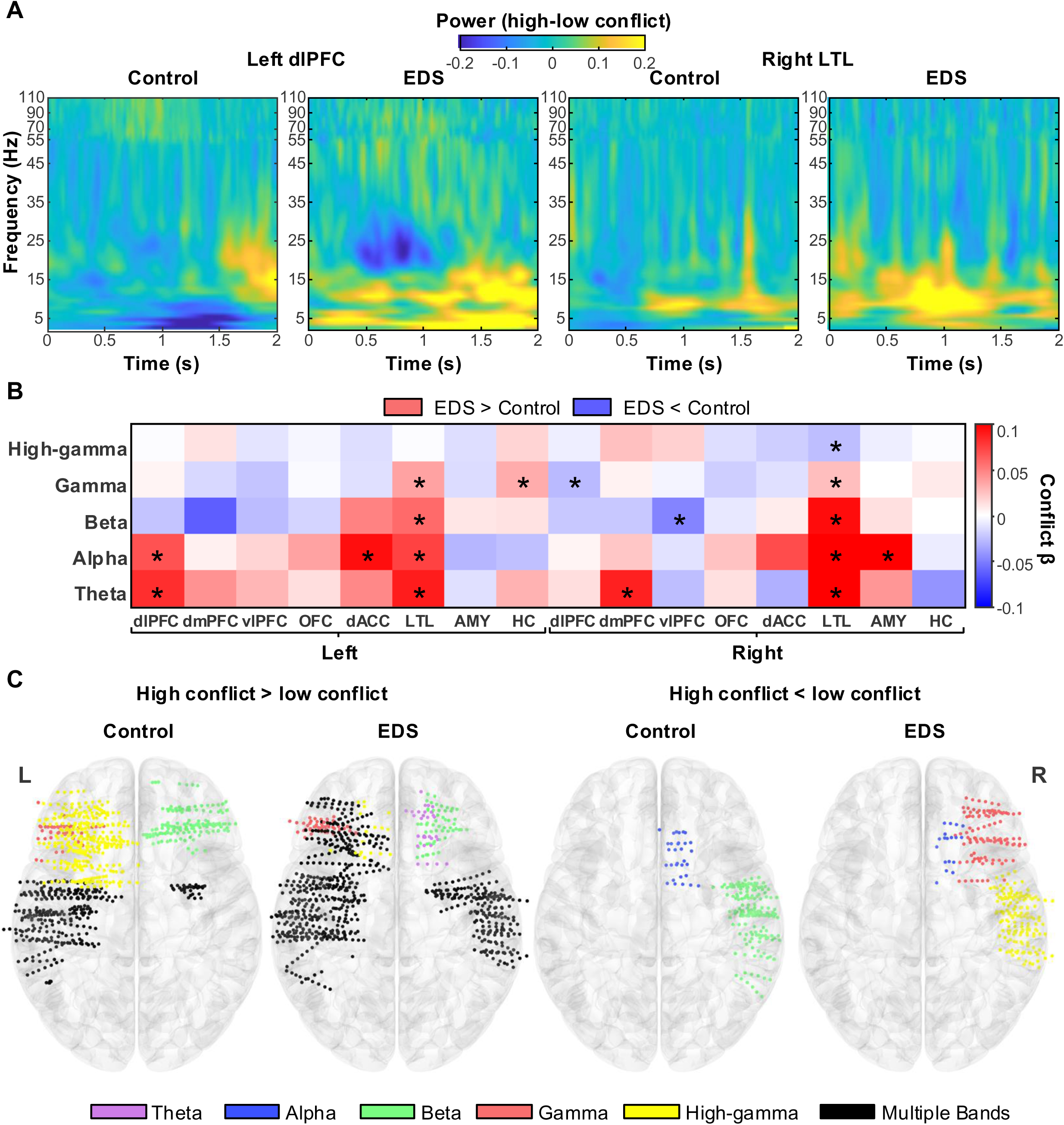
Fronto-temporo-limbic oscillations encoding conflict. **(A)** Time-frequency plots of group-averaged changes in left dlPFC and right LTL power during conflict in control and EDS participants. Intensity values correspond to differences in log-normalized power averaged across channels and participants. **(B)** Heatmap of Conflict*Group predictors in GLMEs of regional band power. Interaction β-weights were coded as the mean difference in conflict-induced change in power (high>low) between groups. Asterisks denote significant interaction predictors (FDR-p<0.05). **(C-E)** Regional electrodes demonstrating significant changes in band power during conflict encoding in control and EDS participants plotted on standard brain templates.

**Table 3.** Effects of Conflict (high>low) on fronto-temporo-limbic power. Summary of post-hoc GLME coefficient contrasts estimating conflict (high>low) effects on spectral power in control and EDS participants. Bolded p-values denote effects that survived FDR-correction.

| Band | Region | Control |  |  | EDS |  |  |
| --- | --- | --- | --- | --- | --- | --- | --- |
| | | $\beta$ | F | p | $\beta$ | F | p |
| Theta | L dlPFC | -0.006 | 0.478 | 0.489 | 0.076 | 44.807 | **<0.001** |
|  | R dmPFC | 0.001 | 0.005 | 0.945 | 0.088 | 13.946 | **<0.001** |
|  | L LTL | 0.071 | 49.174 | **<0.001** | 0.160 | 275.029 | **<0.001** |
|  | R LTL | -0.019 | 4.544 | 0.033 | 0.115 | 109.360 | **<0.001** |
| Alpha | L dlPFC | 0.012 | 1.980 | 0.159 | 0.080 | 51.950 | **<0.001** |
|  | L LTL | 0.085 | 70.950 | **<0.001** | 0.160 | 273.136 | **<0.001** |
|  | R LTL | -0.009 | 0.959 | 0.328 | 0.158 | 202.388 | **<0.001** |
|  | L dACC | -0.018 | 1.218 | 0.270 | 0.077 | 13.887 | **<0.001** |
|  | R AMY | 0.001 | 0.001 | 0.973 | 0.108 | 38.358 | **<0.001** |
| Beta | R vlPFC | 0.045 | 13.246 | **<0.001** | -0.003 | 0.081 | 0.775 |
|  | L LTL | 0.039 | 24.368 | **<0.001** | 0.098 | 167.566 | **<0.001** |
|  | R LTL | -0.021 | 8.855 | **0.003** | 0.074 | 70.135 | **<0.001** |
| Gamma | R dlPFC | 0.008 | 1.925 | 0.165 | -0.019 | 5.875 | **0.015** |
|  | L LTL | 0.001 | 0.066 | 0.798 | 0.038 | 50.163 | **<0.001** |
|  | R LTL | -0.012 | 4.969 | 0.026 | 0.014 | 4.764 | 0.029 |
|  | L HC | 0.002 | 0.062 | 0.804 | 0.034 | 18.437 | **<0.001** |
| High-gamma | R LTL | 0.000 | 0.003 | 0.958 | -0.026 | 37.238 | **<0.001** |

In both groups, conflict increased alpha in left hippocampus alpha, beta in right OFC, gamma/high-gamma in left PFC and dACC, and broadband power in bilateral amygdala, while reducing right dACC alpha power (Fig. 2C, Table S5). Across task conditions, EDS participants showed greater left amygdala beta power, and lower right amygdala theta, left amygdala/hippocampus beta, and right amygdala high-gamma power than controls (Table S5).

#### Fronto-temporo-limbic coherence during conflict encoding in EDS and controls

We then fit GLMEs to pre-response coherence between regions encoding conflict within theta to beta bands (significant effect of conflict on band power in one or both groups) to compare functional connectivity changes encoding conflict between controls and EDS. Conflict x group interactions predicted coherence between region pairs including left dlPFC and hippocampus, right PFC, dACC, and OFC, and bilateral LTL and amygdala (Figs 3A-B, Table S6). In control participants, conflict increased theta/alpha coherence between left dlPFC and left/right LTL, while EDS participants showed reduced left dlPFC-right LTL coherence (Fig. 3C, Table 4).

**Figure 3.**
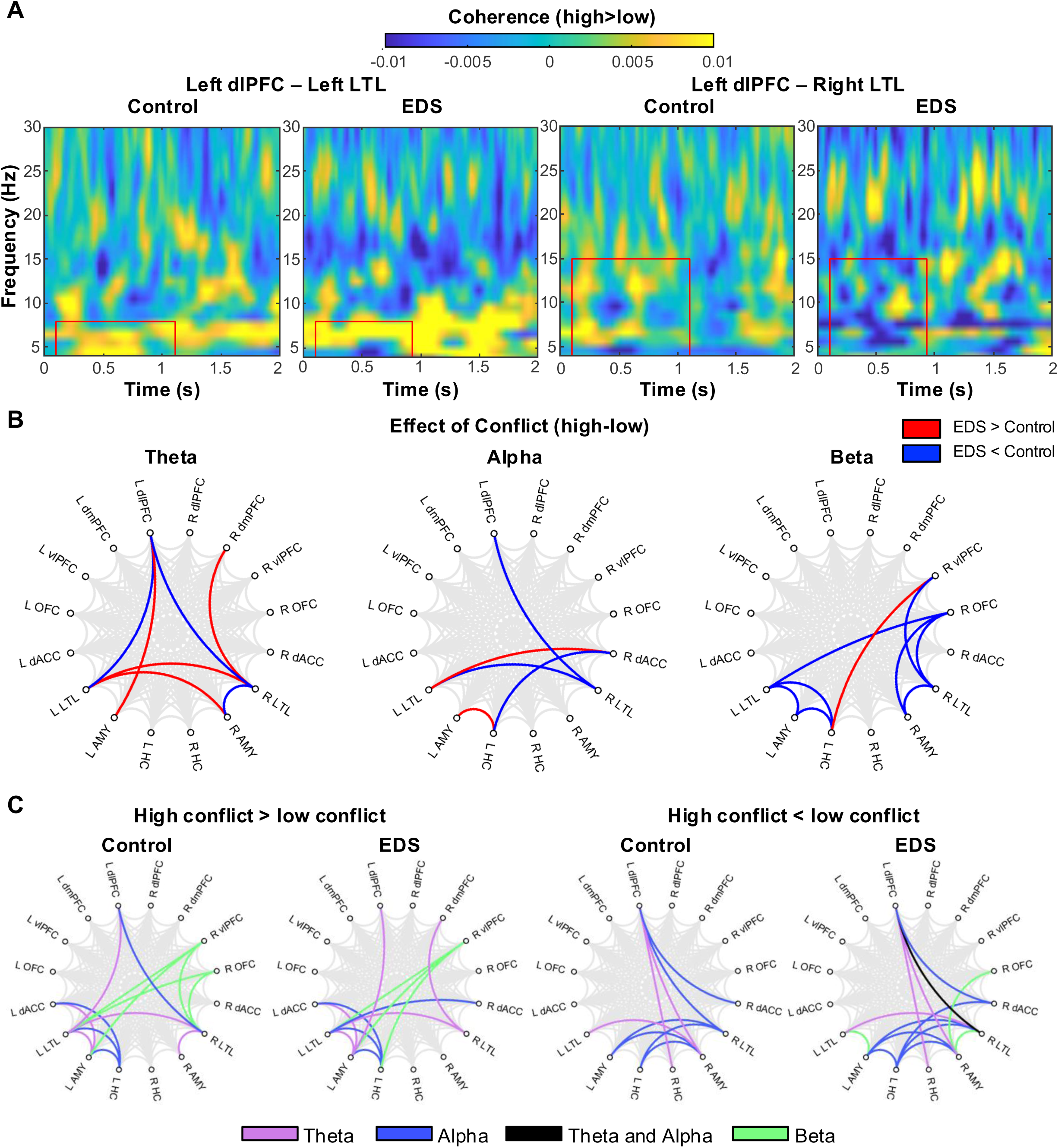
Changes in fronto-temporo-limbic coherence encoding conflict. **(A)** Time-frequency plots of group-averaged changes in left dlPFC-left LTL and left dlPFC-right LTL coherence during conflict encoding in control and EDS participants. Intensity values correspond to differences in coherence or PLV averaged across channel pairs and participants. Inset rectangles correspond to theta activity from 0.1s to median high conflict RT. **(B)** Connectograms depicting significant differences between control and EDS participants in conflict-related coherence (FDR-p<0.05). **(B)** Connectograms depicting significant changes in coherence changes by conflict in each group within theta-beta frequency bands (FDR-p<0.05).

**Table 4.** Effects of Conflict (high>low) on fronto-temporo-limbic coherence. Summary of post-hoc GLME coefficient contrasts of conflict (high>low) effects on coherence in control and EDS participants. Bolded p-values denote effects that survived FDR-correction.

| Band | Connection | Control |  |  | EDS |  |  |
| --- | --- | --- | --- | --- | --- | --- | --- |
| | | $\beta$ | F | p | $\beta$ | F | p |
| Theta |  |  |  |  |  |  |  |
|  | L dlPFC-L LTL | 0.007 | 54.165 | <**0.001** | -0.001 | 2.274 | 0.132 |
|  | L dlPFC-R LTL | 0.001 | 1.311 | 0.252 | -0.006 | 43.135 | <**0.001** |
|  | R dmPFC-R LTL | -0.002 | 1.794 | 0.181 | 0.006 | 6.813 | **0.009** |
|  | L LTL-R LTL | -0.001 | 2.347 | 0.126 | 0.002 | 6.240 | **0.012** |
|  | L dlPFC-L AMY | -0.003 | 3.852 | 0.050 | 0.010 | 18.914 | <**0.001** |
|  | L LTL-R AMY | -0.010 | 22.535 | <**0.001** | 0.002 | 1.641 | 0.200 |
|  | R LTL-R AMY | 0.011 | 22.007 | <**0.001** | -0.005 | 4.640 | 0.031 |
| Alpha |  |  |  |  |  |  |  |
|  | L dlPFC-R LTL | 0.002 | 5.493 | **0.019** | -0.005 | 45.674 | <**0.001** |
|  | L LTL-R LTL | 0.002 | 6.703 | **0.010** | -0.002 | 14.771 | <**0.001** |
|  | L LTL-R dACC | -0.002 | 1.036 | 0.309 | 0.011 | 37.045 | <**0.001** |
|  | R dACC-L HC | 0.000 | 0.000 | 0.998 | -0.014 | 16.865 | <**0.001** |
|  | L AMY-L HC | -0.004 | 1.354 | 0.245 | 0.009 | 6.656 | **0.010** |
| Beta |  |  |  |  |  |  |  |
|  | R OFC-L LTL | 0.004 | 24.126 | <**0.001** | -0.001 | 0.617 | 0.432 |
|  | R OFC-R LTL | 0.003 | 15.789 | <**0.001** | -0.001 | 1.396 | 0.237 |
|  | R vlPFC-R LTL | 0.003 | 5.444 | **0.020** | -0.002 | 5.052 | 0.025 |
|  | L LTL-L AMY | 0.001 | 0.235 | 0.628 | -0.007 | 34.238 | <**0.001** |
|  | R OFC-R AMY | 0.001 | 0.582 | 0.446 | -0.006 | 9.349 | **0.002** |
|  | R LTL-R AMY | 0.001 | 1.491 | 0.222 | -0.004 | 11.029 | **0.001** |
|  | R vlPFC-L HC | 0.000 | 0.102 | 0.750 | 0.005 | 20.549 | <**0.001** |
|  | L LTL-L HC | 0.005 | 34.155 | <**0.001** | 0.001 | 1.474 | 0.225 |
|  | L AMY-L HC | 0.006 | 7.164 | **0.008** | -0.006 | 7.246 | **0.007** |

EDS participants also showed conflict-related increases in right dmPFC-LTL and left dlPFC-amygdala theta, left LTL-right dACC alpha, and right vlPFC-left hippocampus beta coherence. Additionally, conflict reduced left hippocampus-right dACC/left amygdala alpha, left LTL-amygdala beta, and right amygdala-right OFC/LTL beta coherence in EDS. Controls exhibited greater conflict-evoked theta coherence between bilateral LTL-right amygdala, and beta coherence between right OFC-bilateral LTL, right vlPFC-LTL, and left hippocampus-left LTL/amygdala, while theta coherence between bilateral LTL was reduced. Group-level contrasts during high-conflict showed lower coherence in EDS between left dlPFC-bilateral LTL theta/alpha, right dACC-left hippocampus alpha, right OFC-left LTL beta, and left amygdala-hippocampus beta compared to controls (Table S7)

Across participants, conflict increased left LTL-amygdala/hippocampus theta/alpha coherence and right vlPFC-amygdala beta coherence, while reducing theta/alpha coherence between left dlPFC-right amygdala, left dlPFC-right dACC, and right amygdala-left hippocampus. (Figure 3C, Table S8). Alpha coherence between right dACC-amygdala was greater in EDS across task conditions (β=0.018, FDR-p=0.003).

#### dlPFC-dACC-LTL effective connectivity during conflict encoding in EDS and controls

To assess directionality of group differences in fronto-temporo-limbic connectivity during conflict encoding, we used DCM to model baseline-corrected theta-alpha (4-15Hz) cross-spectra between left dlPFC, dACC, LTL, and right LTL in a subset of participants (n=9 control, 7 EDS) with electrode coverage in this network (Fig. 4A). Participants in this subset exhibited similar spectral responses during conflict encoding as our full dataset (Tables S9 & S10). We then used PEB to model group and task (baseline, conflict-encoding) effects on connectivity parameters. We focus here on parameters estimating modulatory effects of conflict on connectivity (high-low) with strong evidence (95% posterior probability) for task/group x task effects.

**Figure 4.**
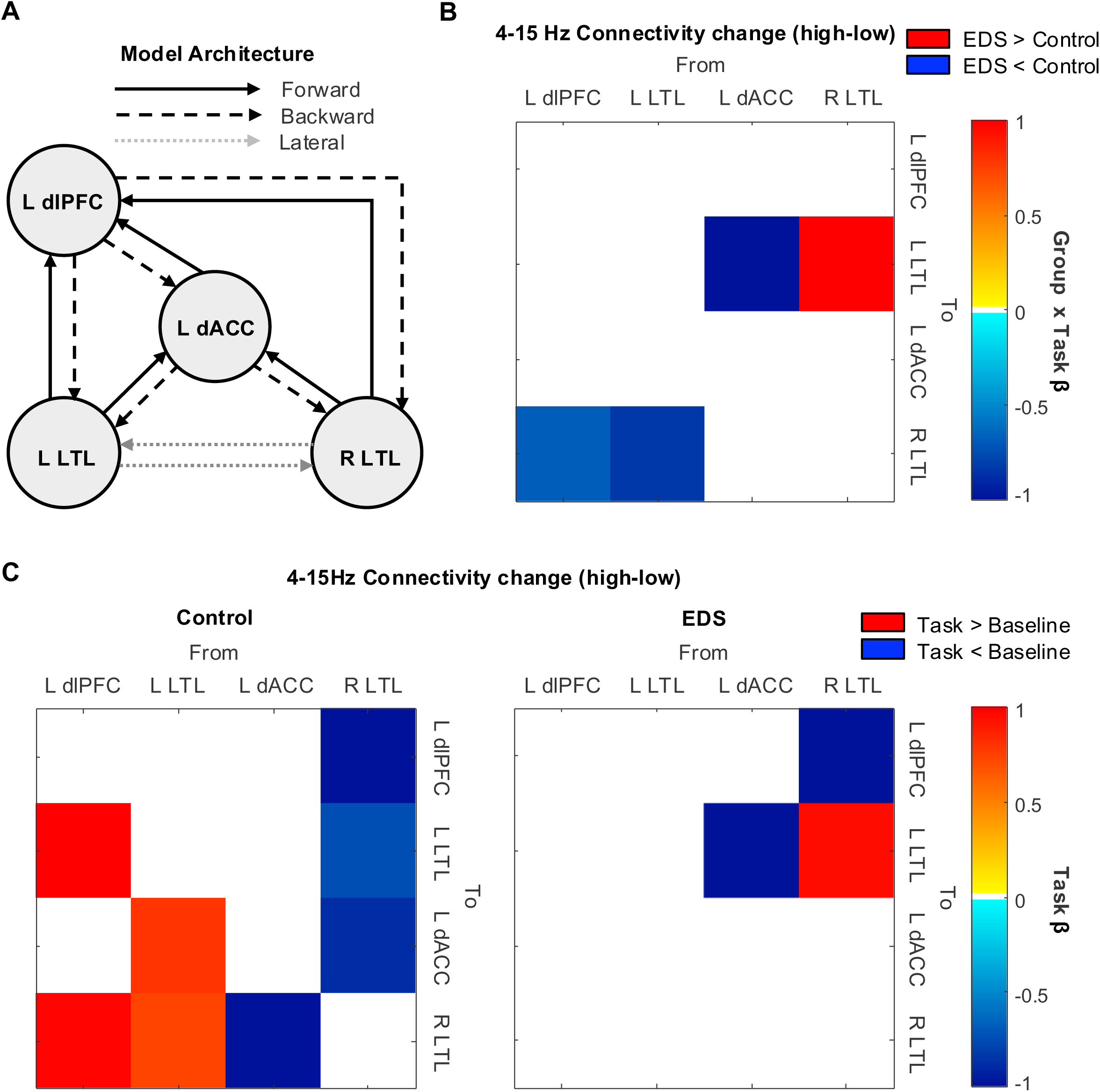
Baseline-corrected model estimates of fronto-temporo-limbic effective connectivity changes encoding conflict. **(A)** Diagram of neural model architecture defined for DCM analyses. (B). PEB model parameters demonstrating strong evidence (95% posterior probability) for group differences in baseline-corrected effective connectivity modulated by conflict (high-low). (C) PEB model parameters demonstrating strong evidence for baseline-corrected effective connectivity modulation by conflict (high-low) in control and EDS participants.

EDS participants showed reduced baseline-corrected effective connectivity from left dlPFC➔right LTL, left dACC➔right LTL, and left LTL➔right LTL, and increased right➔left LTL connectivity relative to controls (Fig. 4B). Post-hoc PEB models found conflict increased baseline-corrected connectivity from left dlPFC➔bilateral LTL, left LTL➔dACC, and left LTL➔right LTL, and reduced connectivity to/from right LTL in controls, while reducing left dACC➔left LTL connectivity in EDS participants (Fig. 4C). Both controls and EDS reduced right LTL➔left dlPFC connectivity during conflict encoding.

## Discussion

We recorded sEEG from fronto-temporo-limbic regions during MSIT performance to examine whether EDS participants encode cognitive conflict differently than controls. Behaviorally, EDS participants exhibited greater conflict-related slowing and responded faster on low-conflict trials, with no difference in accuracy between groups. Spectral analyses showed that conflict elicited widespread fronto-temporo-limbic oscillations in both groups. However, EDS participants exhibited greater theta/alpha responses to conflict in PFC, dACC, LTL, and amygdala, and reduced gamma/high gamma activity in right dlPFC/LTL. Functional connectivity analyses found theta/alpha coherence between left dlPFC and LTL was increased by conflict in controls, but was reduced in EDS. DCM analyses of baseline-corrected theta-alpha effective connectivity found conflict increased left dlPFC➔bilateral LTL connectivity in control, but not EDS participants, while reducing left dACC➔right LTL connectivity in EDS, and right LTL➔left dlPFC connectivity in both groups. In summary, our findings indicate EDS demonstrate heightened oscillatory responses and disrupted fronto-temporo-limbic connectivity during conflict encoding without behavioral impairment in task performance.

### EDS participants demonstrate greater conflict-induced response slowing while maintaining task performance

EDS participants achieved similar accuracy on the MSIT as controls, with greater response time slowing by conflict. Prior cognitive control studies in depression report mixed behavioral findings, with some showing no behavioral impairments,^60-63^ while others report higher error rates and slower response times in depressed individuals.^64-66^ Our findings suggest EDS participants may have needed greater resource allocation over time to maintain task performance. Although it is unclear why EDS participants were faster on low conflict trials, this could reflect anxious/impulsive responding, differences in motor function, or positional factors related to the EMU setting (e.g., keyboard access). Alternatively, cognitive control deficits in EDS, if any, may manifest more robustly at the neural level, rather than in behavioral performance.^37,67^ Therefore, we focus on neural responses during conflict encoding as a more sensitive measure of cognitive function.

### EDS and control participants encode conflict in distributed fronto-temporo-limbic oscillatory networks

In addition to theta and high gamma responses to conflict, we found widespread changes in fronto-temporo-limbic spectral power and coherence encoded conflict in both EDS and controls. Few intracranial studies have directly examined conflict encoding outside of the PFC and dACC,^22,23,68^ as these regions are highly evidenced in cognitive control from scalp EEG and fMRI literature. However, sEEG can capture large-scale neural dynamics with greater spatiotemporal resolution than non-invasive methods. Indeed, intracranial studies have reported conflict encoding oscillations in the OFC,^55,69,70^ amygdala,^71^ and hippocampus.^72,73^ Our findings further evidence multiplexed oscillatory signals encode conflict across distributed fronto-temporo-limbic networks.^74^

### Enhanced fronto-temporo-limbic oscillations in EDS may reflect greater need for cognitive/attentional control

Compared to controls, EDS participants exhibited stronger prefrontal theta-alpha and broadband LTL responses encoding conflict. In both groups, conflict enhanced high-gamma oscillations in left dlPFC, dmPFC, and dACC as in prior sEEG studies of cognitive control.^55,75^ However, only EDS participants showed enhanced theta-alpha responses to conflict in left dlPFC, right dmPFC, and left dACC. These findings may align with previous reports of “hyperfrontality” in depressed individuals when engaging cognitive control.^32,34,35^ Theta-alpha oscillations in dmPFC/dACC are frequently associated with conflict detection, whereas dlPFC responses are thought to serve to top-down cognitive and attention regulation.^6,55,76-78^ Thus, increased prefrontal theta oscillations in EDS participants may reflect greater recruitment of cognitive resources in order to meet task demands^79^ due to inefficient conflict processing.^32^

Notably, differences in conflict encoding between EDS and controls were most pronounced in the LTL. Although not classically linked to cognitive control, the LTL, is sensitive to multiple sources of conflict,^17,54,68,80^ due to its role in visual and semantic processing.^54,81,82^ While both groups exhibited broadband left LTL responses to conflict, only EDS participants demonstrated conflict-related increases in right LTL activity. The right LTL is associated with ventral reorientation of attention to task-relevant stimuli,^83,84^ and right LTL lesions frequently result in spatial neglect.^85^ Heightened broadband responses in right LTL observed in EDS participants may indicate greater recruitment of bottom-up attention/perceptual processing. To our knowledge, LTL activity during conflict tasks has not been widely associated with depressive symptoms. However, one fMRI study found greater inferior temporal lobe activity in depressed participants during response inhibition.^35^ Together, our findings suggest enhanced fronto-temporo-limbic recruitment in EDS may reflect increased effort to maintain task performance during conflict.^84,86^

### Reduced fronto-temporo-limbic connectivity may underlie compensatory oscillatory recruitment in EDS

In controls, conflict enhanced theta–alpha coherence and top-down effective connectivity from left dlPFC➔bilateral LTL, whereas EDS participants reduced coherence and effective connectivity between left dlPFC➔right LTL. Additionally, bottom-up connectivity from right LTL➔left dlPFC was suppressed by conflict in both groups. The LTL modulates stimulus representations to enhance attention towards arousing/salient stimuli,^87-89^ and is thought to be regulated by the dlPFC during increased processing demand.^87^ Enhanced top-down connectivity from left dlPFC➔LTL with reduced bottom-up signaling in controls supports a role for prefrontal regulation of LTL signaling during conflict, which may be disrupted in EDS. Imaging and EEG studies report reduced resting-state functional connectivity of fronto-tempo-limbic circuits in depression,^90,91^ however, the role of these circuits during cognitive control remains unclear. Given the LTL’s role in attention and the involvement of theta-alpha rhythms in control signaling, disrupted dlPFC➔LTL and dACC➔LTL communication in EDS circuits in EDS participants may reflect greater challenge in maintaining task focus.^92,93^ We speculate reduced synchrony and disrupted communication within these circuits in EDS participants may necessitate compensatory increases in oscillatory activity to sustain performance during greater cognitive load.

## Limitations

We address several limitations in interpretation of our findings. While we compared EDS to an epilepsy control group and modeled inter-participant variability, we were unable to fully disentangle effects of depressive symptoms from those related to epilepsy-associated cognitive dysfunction. Studies of conflict encoding in healthy and depressed populations without epilepsy are necessary to determine the generalizability of our study findings. While sEEG studies are mostly limited to epilepsy patients, non-invasive high-density EEG or EEG-fMRI techniques could provide comparable resolution of fronto-temporo-limbic structures. Additionally, heterogeneity in electrode placement and available clinical data limited our ability to distinguish effects of historical versus current depressive symptoms on conflict encoding and assess medication effects. Future work with larger, unmedicated cohorts and standardized diagnostic criteria could clarify whether altered conflict encoding neural features reflect active symptoms, or historical vulnerability for depression.

Additionally, we treated the LTL region as a uniform region due to limitations in electrode coverage. A larger sample could elucidate the role of specific LTL subregions in cognitive control. Finally, while DCM provides biophysical plausibility for models of neural dynamics, our models were limited to the dlPFC, dACC, and LTL by sampling constraints. Future sEEG studies of whole-brain networks, or modulation of these circuits using stimulation paradigms would provide stronger evidence for their role in conflict encoding.

## Conclusion

We provide direct intracranial evidence that cognitive conflict is widely encoded across distributed fronto-temporo-limbic networks. We further show individuals with epilepsy and a history of comorbid depressive symptoms exhibit enhanced theta/alpha fronto-temporo-limbic spectral responses during conflict encoding that are reduced in synchrony and may underlie disrupted top-down regulatory signaling between the dlPFC, dACC, and LTL. These findings suggest that in EDS, heightened fronto-temporo-limbic oscillations during conflict may compensate for disrupted network communication, reflecting greater cognitive recruitment to meet task demands. Our results encourage further studies of fronto-temporo-limbic cognitive control circuits in depression as potential targets for therapeutic intervention.

## Acknowledgements

We gratefully acknowledge technical assistance with data collection from Afsana Afzal, Gavin Belok, Kara Farnes, Madeleine Robertson, Deborah Vallejo-Lopez, and Samuel Zorowitz at MGH. We thank the EMU staff and attending epileptologists and fellows at UCMC for assisting with data collection. We also thank the research participants, whose generosity made this work possible.

Data collection and analysis at UCMC was supported by NIMH R21, #1R21MH127009-01A. Data collection at MGH was supported by grants from the Defense Advanced Research Projects Agency (DARPA) under Cooperative Agreement Number W911NF-14-2-0045 issued by the Army Research Organization (ARO) contracting office in support of DARPA’s SUBNETS Program. The views, opinions, and findings expressed are those of the authors. They should not be interpreted as representing the official views or policies of the Department of Defense, Department of Health & Human Services, any other branch of the U.S. Government, or any other funding entity.

MATLAB code for preprocessing, data analyses, and generating figures is available at https://github.com/ashekara/MSIT-Analysis. Neural and behavioral data supporting the findings of this study will be made available upon request.

## References

1. Administration SAaMHS. Key substance use and mental health indicators in the United States: Results from the 2023 National Survey on Drug Use and Health. 2024. HHS Publication No. PEP24-07-021. https://www.samhsa.gov/data/report/2023-nsduh-annual-national-report

2. Gordon JA. On being a circuit psychiatrist. Nat Neurosci. Oct 26 2016;19(11):1385–1386. doi:10.1038/nn.4419

3. Wexler BE. Returning to basic principles to develop more effective treatments for central nervous system disorders. Exp Biol Med (Maywood). May 2022;247(10):856–867. doi:10.1177/15353702221078291

4. Saveanu R, Etkin A, Duchemin AM, et al. The international Study to Predict Optimized Treatment in Depression (iSPOT-D): outcomes from the acute phase of antidepressant treatment. J Psychiatr Res. Feb 2015;61:1–12. doi:10.1016/j.jpsychires.2014.12.018

5. Cuthbert BN, Insel TR. Toward the future of psychiatric diagnosis: the seven pillars of RDoC. BMC Med. May 14 2013;11(1):126. doi:10.1186/1741-7015-11-126

6. Botvinick MM, Braver TS, Barch DM, Carter CS, Cohen JD. Conflict monitoring and cognitive control. Psychol Rev. Jul 2001;108(3):624-52. doi:10.1037/0033-295x.108.3.624

7. McTeague LM, Huemer J, Carreon DM, Jiang Y, Eickhoff SB, Etkin A. Identification of Common Neural Circuit Disruptions in Cognitive Control Across Psychiatric Disorders. Am J Psychiatry. Jul 1 2017;174(7):676–685. doi:10.1176/appi.ajp.2017.16040400

8. Miller EK, Cohen JD. An integrative theory of prefrontal cortex function. Annu Rev Neurosci. 2001;24(Volume 24, 2001):167-202. doi:10.1146/annurev.neuro.24.1.167

9. Braver TS. The variable nature of cognitive control: a dual mechanisms framework. Trends Cogn Sci. Feb 2012;16(2):106–13. doi:10.1016/j.tics.2011.12.010

10. Yang Z, Oathes DJ, Linn KA, et al. Cognitive Behavioral Therapy Is Associated With Enhanced Cognitive Control Network Activity in Major Depression and Posttraumatic Stress Disorder. Biol Psychiatry Cogn Neurosci Neuroimaging. Apr 2018;3(4):311–319. doi:10.1016/j.bpsc.2017.12.006

11. Widge AS, Heilbronner SR, Hayden BY. Prefrontal cortex and cognitive control: new insights from human electrophysiology. F1000Res. 2019;8doi:10.12688/f1000research.20044.1

12. Stroop JR. Studies of interference in serial verbal reactions. Journal of experimental psychology. 1935;18(6):643.

13. Kim C, Kroger JK, Kim J. A functional dissociation of conflict processing within anterior cingulate cortex. Hum Brain Mapp. Feb 2011;32(2):304–12. doi:10.1002/hbm.21020

14. Boschin EA, Brkic MM, Simons JS, Buckley MJ. Distinct Roles for the Anterior Cingulate and Dorsolateral Prefrontal Cortices During Conflict Between Abstract Rules. Cereb Cortex. Jan 1 2017;27(1):34–45. doi:10.1093/cercor/bhw350

15. Cavanagh JF, Frank MJ. Frontal theta as a mechanism for cognitive control. Trends Cogn Sci. Aug 2014;18(8):414–21. doi:10.1016/j.tics.2014.04.012

16. Myers JC, Chinn LK, Sur S, Golob EJ. Widespread theta coherence during spatial cognitive control. Neuropsychologia. Sep 17 2021;160:107979. doi:10.1016/j.neuropsychologia.2021.107979

17. Bush G, Shin LM. The Multi-Source Interference Task: an fMRI task that reliably activates the cingulo-frontal-parietal cognitive/attention network. Nat Protoc. 2006;1(1):308–13. doi:10.1038/nprot.2006.48

18. Robertson JA, Thomas AW, Prato FS, Johansson M, Nittby H. Simultaneous fMRI and EEG during the multi-source interference task. PLoS One. 2014;9(12):e114599. doi:10.1371/journal.pone.0114599

19. Gonzalez-Villar AJ, Carrillo-de-la-Pena MT. Brain electrical activity signatures during performance of the Multisource Interference Task. Psychophysiology. Jun 2017;54(6):874–881. doi:10.1111/psyp.12843

20. Provenza NR, Paulk AC, Peled N, et al. Decoding task engagement from distributed network electrophysiology in humans. J Neural Eng. Aug 16 2019;16(5):056015. doi:10.1088/1741-2552/ab2c58

21. Avvaru S, Peled N, Provenza NR, Widge AS, Parhi KK. Region-Level Functional and Effective Network Analysis of Human Brain During Cognitive Task Engagement. IEEE Trans Neural Syst Rehabil Eng. 2021;29:1651–1660. doi:10.1109/TNSRE.2021.3105432

22. Basu I, Yousefi A, Crocker B, et al. Closed-loop enhancement and neural decoding of cognitive control in humans. Nat Biomed Eng. Apr 2023;7(4):576–588. doi:10.1038/s41551-021-00804-y

23. Haegens S, Pathak YJ, Smith EH, et al. Alpha and broadband high-frequency activity track task dynamics and predict performance in controlled decision-making. Psychophysiology. May 2022;59(5):e13901. doi:10.1111/psyp.13901

24. Karnath HO. New insights into the functions of the superior temporal cortex. Nat Rev Neurosci. Aug 2001;2(8):568–76. doi:10.1038/35086057

25. Zatorre RJ, Belin P. Spectral and temporal processing in human auditory cortex. Cereb Cortex. Oct 2001;11(10):946–53. doi:10.1093/cercor/11.10.946

26. Visser M, Jefferies E, Lambon Ralph MA. Semantic processing in the anterior temporal lobes: a meta-analysis of the functional neuroimaging literature. J Cogn Neurosci. Jun 2010;22(6):1083–94. doi:10.1162/jocn.2009.21309

27. Collins JA, Olson IR. Beyond the FFA: The role of the ventral anterior temporal lobes in face processing. Neuropsychologia. Aug 2014;61:65–79. doi:10.1016/j.neuropsychologia.2014.06.005

28. Leonard MK, Chang EF. Dynamic speech representations in the human temporal lobe. Trends Cogn Sci. Sep 2014;18(9):472–9. doi:10.1016/j.tics.2014.05.001

29. Ren Y, Pan L, Du X, Hou Y, Li X, Song Y. Functional brain network mechanism of executive control dysfunction in temporal lobe epilepsy. BMC Neurol. Apr 15 2020;20(1):137. doi:10.1186/s12883-020-01711-6

30. Harden CL. The co-morbidity of depression and epilepsy: epidemiology, etiology, and treatment. Neurology. Sep 24 2002;59(6 Suppl 4):S48-55. doi:10.1212/wnl.59.6_suppl_4.s48

31. Piazzini A, Canevini MP, Maggiori G, Canger R. Depression and Anxiety in Patients with Epilepsy. Epilepsy Behav. Oct 2001;2(5):481–489. doi:10.1006/ebeh.2001.0247

32. Wagner G, Sinsel E, Sobanski T, et al. Cortical inefficiency in patients with unipolar depression: an event-related FMRI study with the Stroop task. Biol Psychiatry. May 15 2006;59(10):958–65. doi:10.1016/j.biopsych.2005.10.025

33. Grahek I, Everaert J, Krebs RM, Koster EHW. Cognitive Control in Depression: Toward Clinical Models Informed by Cognitive Neuroscience. Clinical Psychological Science. 2018;6(4):464–480. doi:10.1177/2167702618758969

34. Krompinger JW, Simons RF. Cognitive inefficiency in depressive undergraduates: stroop processing and ERPs. Biol Psychol. Mar 2011;86(3):239–46. doi:10.1016/j.biopsycho.2010.12.004

35. Langenecker SA, Kennedy SE, Guidotti LM, et al. Frontal and limbic activation during inhibitory control predicts treatment response in major depressive disorder. Biol Psychiatry. Dec 1 2007;62(11):1272–80. doi:10.1016/j.biopsych.2007.02.019

36. Holmes AJ, Pizzagalli DA. Response conflict and frontocingulate dysfunction in unmedicated participants with major depression. Neuropsychologia. Oct 2008;46(12):2904–13. doi:10.1016/j.neuropsychologia.2008.05.028

37. Paulus MP. Cognitive control in depression and anxiety: out of control? Current Opinion in Behavioral Sciences. 2015;1:113–120. doi:10.1016/j.cobeha.2014.12.003

38. Beck AT, Ward CH, Mendelson M, Mock J, Erbaugh J. An inventory for measuring depression. Arch Gen Psychiatry. Jun 1961;4(6):561–71. doi:10.1001/archpsyc.1961.01710120031004

39. Kroenke K, Spitzer RL, Williams JB. The PHQ-9: validity of a brief depression severity measure. J Gen Intern Med. Sep 2001;16(9):606–13. doi:10.1046/j.1525-1497.2001.016009606.x

40. Bush G, Shin LM, Holmes J, Rosen BR, Vogt BA. The Multi-Source Interference Task: validation study with fMRI in individual subjects. Mol Psychiatry. Jan 2003;8(1):60–70. doi:10.1038/sj.mp.4001217

41. Soper DJ, Reich D, Ross A, et al. Modular pipeline for reconstruction and localization of implanted intracranial ECoG and sEEG electrodes. PLoS One. 2023;18(7):e0287921. doi:10.1371/journal.pone.0287921

42. Fischl B. FreeSurfer. Neuroimage. Aug 15 2012;62(2):774-81. doi:10.1016/j.neuroimage.2012.01.021

43. Klein A, Tourville J. 101 labeled brain images and a consistent human cortical labeling protocol. Front Neurosci. 2012;6:171. doi:10.3389/fnins.2012.00171

44. Felsenstein O, Peled N, Hahn E, et al. Multi-modal neuroimaging analysis and visualization tool (MMVT). arXiv preprint arXiv:191210079. 2019;

45. Rosenberg JR, Amjad AM, Breeze P, Brillinger DR, Halliday DM. The Fourier approach to the identification of functional coupling between neuronal spike trains. Prog Biophys Mol Biol. 1989;53(1):1–31. doi:10.1016/0079-6107(89)90004-7

46. Brillinger DR. Time series: data analysis and theory. SIAM; 2001.

47. Friston KJ, Harrison L, Penny W. Dynamic causal modelling. Neuroimage. Aug 2003;19(4):1273-302. doi:10.1016/s1053-8119(03)00202-7

48. Cohen MX, Cavanagh JF. Single-trial regression elucidates the role of prefrontal theta oscillations in response conflict. Front Psychol. 2011;2:30. doi:10.3389/fpsyg.2011.00030

49. Pinotsis DA, Fitzgerald S, See C, Sementsova A, Widge AS. Toward biophysical markers of depression vulnerability. Front Psychiatry. 2022;13:938694. doi:10.3389/fpsyt.2022.938694

50. Sheth SA, Mian MK, Patel SR, et al. Human dorsal anterior cingulate cortex neurons mediate ongoing behavioural adaptation. Nature. Aug 9 2012;488(7410):218–21. doi:10.1038/nature11239

51. Penny WD, Friston KJ, Ashburner JT, Kiebel SJ, Nichols TE. Statistical parametric mapping: the analysis of functional brain images. Elsevier; 2011.

52. Friston KJ, Bastos A, Litvak V, Stephan KE, Fries P, Moran RJ. DCM for complex-valued data: cross-spectra, coherence and phase-delays. Neuroimage. Jan 2 2012;59(1):439–55. doi:10.1016/j.neuroimage.2011.07.048

53. Moran RJ, Kiebel SJ, Stephan KE, Reilly RB, Daunizeau J, Friston KJ. A neural mass model of spectral responses in electrophysiology. Neuroimage. Sep 1 2007;37(3):706–20. doi:10.1016/j.neuroimage.2007.05.032

54. Niendam TA, Laird AR, Ray KL, Dean YM, Glahn DC, Carter CS. Meta-analytic evidence for a superordinate cognitive control network subserving diverse executive functions. Cogn Affect Behav Neurosci. Jun 2012;12(2):241–68. doi:10.3758/s13415-011-0083-5

55. Oehrn CR, Hanslmayr S, Fell J, et al. Neural communication patterns underlying conflict detection, resolution, and adaptation. J Neurosci. Jul 30 2014;34(31):10438–52. doi:10.1523/JNEUROSCI.3099-13.2014

56. Marreiros AC, Cagnan H, Moran RJ, Friston KJ, Brown P. Basal ganglia-cortical interactions in Parkinsonian patients. Neuroimage. Feb 1 2013;66:301–10. doi:10.1016/j.neuroimage.2012.10.088

57. fdr_bh. MATLAB Central File Exchange; 2024. https://www.mathworks.com/matlabcentral/fileexchange/27418-fdr_bh

58. Friston KJ, Litvak V, Oswal A, et al. Bayesian model reduction and empirical Bayes for group (DCM) studies. Neuroimage. Mar 2016;128:413–431. doi:10.1016/j.neuroimage.2015.11.015

59. Zeidman P, Jafarian A, Seghier ML, et al. A guide to group effective connectivity analysis, part 2: Second level analysis with PEB. Neuroimage. Oct 15 2019;200:12–25. doi:10.1016/j.neuroimage.2019.06.032

60. Vanderhasselt MA, De Raedt R. Impairments in cognitive control persist during remission from depression and are related to the number of past episodes: an event related potentials study. Biol Psychol. Jul 2009;81(3):169–76. doi:10.1016/j.biopsycho.2009.03.009

61. Holmes AJ, Pizzagalli DA. Effects of task-relevant incentives on the electrophysiological correlates of error processing in major depressive disorder. Cogn Affect Behav Neurosci. Mar 2010;10(1):119–28. doi:10.3758/CABN.10.1.119

62. Meiran N, Diamond GM, Toder D, Nemets B. Cognitive rigidity in unipolar depression and obsessive compulsive disorder: examination of task switching, Stroop, working memory updating and post-conflict adaptation. Psychiatry Res. Jan 30 2011;185(1-2):149–56. doi:10.1016/j.psychres.2010.04.044

63. Davey CG, Yucel M, Allen NB, Harrison BJ. Task-related deactivation and functional connectivity of the subgenual cingulate cortex in major depressive disorder. Front Psychiatry. 2012;3:14. doi:10.3389/fpsyt.2012.00014

64. Ravnkilde B, Videbech P, Clemmensen K, Egander A, Rasmussen NA, Rosenberg R. Cognitive deficits in major depression. Scand J Psychol. Jul 2002;43(3):239–51. doi:10.1111/1467-9450.00292

65. Gohier B, Ferracci L, Surguladze SA, et al. Cognitive inhibition and working memory in unipolar depression. J Affect Disord. Jul 2009;116(1-2):100–5. doi:10.1016/j.jad.2008.10.028

66. Snyder HR. Major depressive disorder is associated with broad impairments on neuropsychological measures of executive function: a meta-analysis and review. Psychol Bull. Jan 2013;139(1):81–132. doi:10.1037/a0028727

67. Ng J, Chan HY, Schlaghecken F. Dissociating effects of subclinical anxiety and depression on cognitive control. Adv Cogn Psychol. 2012;8(1):38–49. doi:10.2478/v10053-008-0100-6

68. Xiao Y, Chou CC, Cosgrove GR, et al. Cross-task specificity and within-task invariance of cognitive control processes. Cell Rep. Jan 31 2023;42(1):111919. doi:10.1016/j.celrep.2022.111919

69. Tang H, Yu HY, Chou CC, et al. Cascade of neural processing orchestrates cognitive control in human frontal cortex. Elife. Feb 18 2016;5doi:10.7554/eLife.12352

70. Chen KH, Tang AM, Gilbert ZD, et al. Theta low-gamma phase amplitude coupling in the human orbitofrontal cortex increases during a conflict-processing task. J Neural Eng. Feb 16 2022;19(1)doi:10.1088/1741-2552/ac4f9b

71. Tang AM, Chen KH, Gogia AS, et al. Amygdaloid theta-band power increases during conflict processing in humans. J Clin Neurosci. Sep 2021;91:183–192. doi:10.1016/j.jocn.2021.07.001

72. Chen KH, Gogia AS, Tang AM, et al. Beta-band modulation in the human hippocampus during a conflict response task. J Neural Eng. Nov 11 2020;17(6)doi:10.1088/1741-2552/abc1b8

73. Oehrn CR, Baumann C, Fell J, et al. Human Hippocampal Dynamics during Response Conflict. Curr Biol. Aug 31 2015;25(17):2307–13. doi:10.1016/j.cub.2015.07.032

74. Helfrich RF, Knight RT. Oscillatory Dynamics of Prefrontal Cognitive Control. Trends Cogn Sci. Dec 2016;20(12):916–930. doi:10.1016/j.tics.2016.09.007

75. Bartoli E, Conner CR, Kadipasaoglu CM, et al. Temporal Dynamics of Human Frontal and Cingulate Neural Activity During Conflict and Cognitive Control. Cereb Cortex. Nov 1 2018;28(11):3842–3856. doi:10.1093/cercor/bhx245

76. MacDonald AW, 3rd, Cohen JD, Stenger VA, Carter CS. Dissociating the role of the dorsolateral prefrontal and anterior cingulate cortex in cognitive control. Science. Jun 9 2000;288(5472):1835–8. doi:10.1126/science.288.5472.1835

77. Hanslmayr S, Pastotter B, Bauml KH, Gruber S, Wimber M, Klimesch W. The electrophysiological dynamics of interference during the Stroop task. J Cogn Neurosci. Feb 2008;20(2):215–25. doi:10.1162/jocn.2008.20020

78. Hopfinger JB, Buonocore MH, Mangun GR. The neural mechanisms of top-down attentional control. Nat Neurosci. Mar 2000;3(3):284–91. doi:10.1038/72999

79. Sadaghiani S, Kleinschmidt A. Brain Networks and alpha-Oscillations: Structural and Functional Foundations of Cognitive Control. Trends Cogn Sci. Nov 2016;20(11):805–817. doi:10.1016/j.tics.2016.09.004

80. Fan J, Flombaum JI, McCandliss BD, Thomas KM, Posner MI. Cognitive and brain consequences of conflict. Neuroimage. Jan 2003;18(1):42–57. doi:10.1006/nimg.2002.1319

81. Banich MT, Milham MP, Jacobson BL, et al. Attentional selection and the processing of task-irrelevant information: insights from fMRI examinations of the Stroop task. Prog Brain Res. 2001;134:459–70. doi:10.1016/s0079-6123(01)34030-x

82. Milham MP, Erickson KI, Banich MT, et al. Attentional control in the aging brain: insights from an fMRI study of the stroop task. Brain Cogn. Aug 2002;49(3):277–96. doi:10.1006/brcg.2001.1501

83. Fox MD, Corbetta M, Snyder AZ, Vincent JL, Raichle ME. Spontaneous neuronal activity distinguishes human dorsal and ventral attention systems. Proc Natl Acad Sci U S A. Jun 27 2006;103(26):10046–51. doi:10.1073/pnas.0604187103

84. Corbetta M, Shulman GL. Control of goal-directed and stimulus-driven attention in the brain. Nat Rev Neurosci. Mar 2002;3(3):201–15. doi:10.1038/nrn755

85. Hillis AE, Newhart M, Heidler J, Barker PB, Herskovits EH, Degaonkar M. Anatomy of spatial attention: insights from perfusion imaging and hemispatial neglect in acute stroke. J Neurosci. Mar 23 2005;25(12):3161–7. doi:10.1523/JNEUROSCI.4468-04.2005

86. Stemmann H, Freiwald WA. Evidence for an attentional priority map in inferotemporal cortex. Proc Natl Acad Sci U S A. Nov 19 2019;116(47):23797–23805. doi:10.1073/pnas.1821866116

87. Mitchell DG, Nakic M, Fridberg D, Kamel N, Pine DS, Blair RJ. The impact of processing load on emotion. Neuroimage. Feb 1 2007;34(3):1299–309. doi:10.1016/j.neuroimage.2006.10.012

88. Pessoa L, Kastner S, Ungerleider LG. Attentional control of the processing of neural and emotional stimuli. Brain Res Cogn Brain Res. Dec 2002;15(1):31–45. doi:10.1016/s0926-6410(02)00214-8

89. Herzog AG, Van Hoesen GW. Temporal neocortical afferent connections to the amygdala in the rhesus monkey. Brain Res. Oct 8 1976;115(1):57–69. doi:10.1016/0006-8993(76)90822-2

90. McVoy M, Aebi ME, Loparo K, et al. Resting-State Quantitative Electroencephalography Demonstrates Differential Connectivity in Adolescents with Major Depressive Disorder. J Child Adolesc Psychopharmacol. Jun 2019;29(5):370–377. doi:10.1089/cap.2018.0166

91. Rolls ET, Cheng W, Gong W, et al. Functional Connectivity of the Anterior Cingulate Cortex in Depression and in Health. Cereb Cortex. Jul 22 2019;29(8):3617–3630. doi:10.1093/cercor/bhy236

92. Keller AS, Leikauf JE, Holt-Gosselin B, Staveland BR, Williams LM. Paying attention to attention in depression. Transl Psychiatry. Nov 7 2019;9(1):279. doi:10.1038/s41398-019-0616-1

93. Keller AS, Payne L, Sekuler R. Characterizing the roles of alpha and theta oscillations in multisensory attention. Neuropsychologia. May 2017;99:48–63. doi:10.1016/j.neuropsychologia.2017.02.021

