## Supplemental Methods, Table S1, Table S4, Table S7 for "Individuals with epilepsy and a history of comorbid depressive symptoms exhibit disrupted fronto-temporo-limbic connectivity during cognitive conflict encoding"

**sEEG acquisition**

sEEG data were recorded from stereotactic depth electrodes (Ad-Tech Medical or PMT) that were 0.8-1.0mm in diameter with 8-16 platinum/iridium contacts 1-2.4mm in length. At the time of acquisition, depth recordings were referenced to an EEG electrode placed on the skin at either cervical vertebra 2 or Cz. Image onset times were synchronized with sEEG data using a transistor-transistor logic (TTL) trigger generated by a PCI parallel port output from MATLAB (MGH/BWH), or a photodiode placed on the bottom-right of the laptop screen (UCMC). For the latter, when a response was made, a bright circle undetectable to the participant would appear on the laptop screen at the photodiode’s position. Analog photodiode voltages and TTL signals during the task were recorded by the EEG acquisition system.

**sEEG preprocessing**

sEEG recordings were pre-processed using Fieldtrip^1^ and custom MATLAB scripts. Channel data from sEEGs were aligned with stimulus onset times and epoched to 5s stimulus-locked trials from -2 to +3s with respect to stimulus onset. We included 1s of buffer data at the ends of each trial to account for edge effects induced by Morlet wavelet transformation. Channel data were then bipolar re-referenced, high-pass filtered at 0.5Hz (5^th^-order Butterworth filter) if low-frequency artifacts were visually observed, notch-filtered between 55-65Hz (4^th^-order Butterworth filter) to remove line noise, and then down-sampled to 512Hz to account for differences in sampling rate. Inter-ictal spike activity and amplifier saturation artifacts were labeled by a z-score threshold set for each participant and manually inspected. Trials with labeled artifacts were excluded from neural analyses.

**Canonical time-frequency analysis**

We estimated stimulus-locked time-frequency power in theta (4-8Hz), alpha (8-15Hz), beta (15-30Hz), gamma (30-55Hz), and high gamma (70-110Hz) bands using Morlet wavelet transformation in Fieldtrip and custom MATLAB scripts. Wavelets were set to 7 cycles and a width of 3 SD, and spectral decomposition was performed with time resolution of 23.4ms and frequency resolution of 1Hz (4-55Hz) or 5Hz (70-110Hz).

Spectral coherence^2^ was used to measure amplitude and phase synchronization of frontotemporal oscillations between 4-30Hz with time resolution as above and frequency resolution of 1Hz:

$Coherence_{xy}(t,\omega)= \frac{|S_{xy}(t,\omega)|}{\sqrt{S_{xx}{\left( t,\omega\right)S}_{yy}(t,\omega)}}$	(1)

where for each time-frequency point $(t,\omega)$, $S_{xy}$ denotes the trial-averaged cross spectral density of signals $x$ and $y$, and $\boldsymbol{S}_{\boldsymbol{xx}}$ and $\boldsymbol{S}_{\boldsymbol{yy}}$ denote their respective trial-averaged power spectral densities. Coherence estimates were performed separately for low conflict and high conflict trials.

**Statistical Analysis: Selection of data transformation and GLME parameters**

We estimated distributions of behavioral and neural data (*allfitdist*^3^) and selected fitted distributions with the lowest Akaike information criterion (AIC) to select GLME distributions and link functions. GLMEs were then fit to raw or transformed response variables using a link function suitable for the approximated data distribution. Model performance was assessed by visually inspecting Q-Q plots of residual normality. For analysis of behavioral and neural response data, log-transformed response times, and baseline-normalized power and coherence data all approximated normal distributions. GLMEs of behavioral and neural data using an identity link had approximately normally residuals and yielded the best performance.

**Table S1. DKT40 atlas definitions of regions selected for analysis.** Atlas labels were combined for prefrontal, temporal, and orbitofrontal regions of interest. Electrodes localized to atlas labels for a given region were pooled for region-specific analyses.

| **Region** | **DKT40 labels** |
| --- | --- |
| Left dlPFC | left rostral middle frontal, left caudal middle frontal |
| Right dlPFC | right rostral middle frontal, right caudal middle frontal |
| Left dmPFC | left superior frontal |
| Right dmPFC | right superior frontal |
| Left vlPFC | left pars triangularis, left pars orbitalis |
| Right vlPFC | right pars triangularis, right pars orbitalis |
| Left OFC | left medial orbitofrontal, left lateral orbitofrontal |
| Right OFC | right medial orbitofrontal, right lateral orbitofrontal |
| Left LTL | left superior temporal, left middle temporal, left inferior temporal, left transverse temporal |
| Right LTL | right superior temporal, right middle temporal, right inferior temporal, right transverse temporal |
| Left dACC | left caudal anterior cingulate |
| Right dACC | right caudal anterior cingulate |
| Left amygdala | left amygdala |
| Right amygdala | right amygdala |
| Left hippocampus | left hippocampus |
| Right hippocampus | right hippocampus |

**Table S2. Demographic and clinical characteristics of MGH/BWH and UCMC participants.** Seizure foci were determined based on clinical consensus following the monitoring period. Psychiatric histories and BDI-II or PHQ-9 scores of MGH participants, where available, were obtained from neuropsychiatric evaluations prior to implantation procedures. Psychiatric histories were obtained from neurobehavioral interviews performed prior to or during monitoring, or from chart reviews. PHQ-9 scores of UCMC participants, where available, were obtained during the monitoring period.

**Table S3. Summary of Conflict x Group GLMEs of fronto-temporo-limbic power.** P-values were FDR-corrected for 80 models of band power with a critical value of p=0.006. Bolded p-values denote effects that survived FDR-correction.

**Table S4. Effects of Group (EDS>control) on fronto-temporo-limbic power.** Summary of post-hoc GLME coefficient contrasts estimating group (high>low) differences in spectral power during low and high conflict trials. Bolded p-values denote effects that survived FDR-correction.

|  |  | **Low Conflict** | | | **High conflict** | | |
| --- | --- | --- | --- | --- | --- | --- | --- |
| **Band** | **Region** | **β** | **F** | **p** | **β** | **F** | **p** |
| Theta |  |  |  |  |  |  |  |
|  | L dlPFC | -0.060 | 0.564 | 0.453 | 0.022 | 0.076 | 0.783 |
|  | R dmPFC | -0.077 | 0.991 | 0.320 | 0.010 | 0.015 | 0.903 |
|  | L LTL | 0.057 | 0.299 | 0.585 | 0.145 | 1.963 | 0.161 |
|  | R LTL | -0.098 | 1.976 | 0.160 | 0.036 | 0.273 | 0.602 |
| Alpha |  |  |  |  |  |  |  |
|  | L dlPFC | -0.083 | 1.540 | 0.215 | -0.016 | 0.053 | 0.817 |
|  | L LTL | 0.064 | 0.591 | 0.442 | 0.138 | 2.737 | 0.098 |
|  | R LTL | -0.065 | 0.973 | 0.324 | 0.102 | 2.428 | 0.119 |
|  | L dACC | 0.002 | 0.001 | 0.979 | 0.097 | 1.180 | 0.277 |
|  | R AMY | -0.052 | 0.956 | 0.328 | 0.055 | 1.063 | 0.303 |
| Beta |  |  |  |  |  |  |  |
|  | R vlPFC | -0.057 | 0.777 | 0.378 | -0.105 | 2.615 | 0.106 |
|  | L LTL | -0.005 | 0.007 | 0.934 | 0.053 | 0.699 | 0.403 |
|  | R LTL | -0.038 | 0.647 | 0.421 | 0.056 | 1.388 | 0.239 |
| Gamma |  |  |  |  |  |  |  |
|  | R dlPFC | -0.012 | 0.085 | 0.770 | -0.039 | 0.866 | 0.352 |
|  | L LTL | -0.017 | 0.413 | 0.520 | 0.019 | 0.506 | 0.477 |
|  | R LTL | 0.014 | 0.225 | 0.635 | 0.040 | 1.733 | 0.188 |
|  | L HC | -0.008 | 1.061 | 0.303 | 0.024 | 9.568 | **0.002** |
| High-gamma | |  |  |  |  |  |  |
|  | R LTL | 0.004 | 0.038 | 0.846 | -0.022 | 1.232 | 0.267 |

**Table S5. Summary of reduced GLMEs of fronto-temporo-limbic power.** P-values were FDR-corrected for 64 reduced models of band power with a critical value of p=0.007. Bolded p-values denote effects that survived FDR-correction.

**Table S6. Summary of Conflict x Group GLMEs of fronto-temporo-limbic coherence.** P-values were FDR-corrected for 64 coherence models with a critical value of p = 0.008. Bolded p-values denote effects that survived FDR-correction.

**Table S7. Effects of Group (EDS>control) on fronto-temporo-limbic coherence.** Summary of post-hoc GLME coefficient contrasts estimating group (high>low) differences in spectral coherence during low and high conflict trials. Bolded p-values denote effects that survived FDR-correction.

|  |  | **EDS>Control Low Conflict** | | | **EDS>Control High conflict** | | |
| --- | --- | --- | --- | --- | --- | --- | --- |
| **Band** | **Region** | **β** | **F** | **p** | **β** | **F** | **p** |
| Theta |  |  |  |  |  |  |  |
|  | L dlPFC-L LTL | -0.001 | 0.202 | 0.653 | -0.010 | 8.746 | **0.003** |
|  | L dlPFC-R LTL | 0.006 | 2.811 | 0.094 | -0.001 | 0.036 | 0.850 |
|  | R dmPFC-R LTL | -0.005 | 1.848 | 0.174 | 0.003 | 0.655 | 0.418 |
|  | L LTL-R LTL | 0.001 | 0.057 | 0.811 | 0.004 | 1.224 | 0.269 |
|  | L dlPFC-L AMY | -0.009 | 6.150 | **0.013** | 0.004 | 1.131 | 0.288 |
|  | L LTL-R AMY | -0.004 | 0.846 | 0.358 | 0.008 | 2.782 | 0.095 |
|  | R LTL-R AMY | 0.010 | 1.195 | 0.274 | -0.007 | 0.550 | 0.458 |
| Alpha |  |  |  |  |  |  |  |
|  | L dlPFC-R LTL | 0.002 | 2.216 | 0.137 | -0.004 | 8.347 | **0.004** |
|  | L LTL-R LTL | 0.005 | 6.509 | **0.011** | 0.000 | 0.017 | 0.896 |
|  | L LTL-R dACC | -0.007 | 5.963 | **0.015** | 0.006 | 4.333 | 0.038 |
|  | R dACC-L HC | 0.005 | 1.509 | 0.220 | -0.009 | 6.391 | **0.012** |
|  | L AMY-L HC | 0.007 | 0.829 | 0.363 | 0.021 | 6.349 | **0.012** |
| Beta |  |  |  |  |  |  |  |
|  | R OFC-L LTL | 0.001 | 1.402 | 0.237 | -0.003 | 5.928 | **0.015** |
|  | R OFC-R LTL | 0.002 | 0.561 | 0.454 | -0.002 | 0.618 | 0.432 |
|  | R vlPFC-R LTL | 0.003 | 1.219 | 0.270 | -0.001 | 0.103 | 0.748 |
|  | L LTL-L AMY | 0.002 | 0.616 | 0.432 | -0.005 | 2.485 | 0.115 |
|  | R OFC-R AMY | 0.006 | 3.049 | 0.081 | -0.002 | 0.329 | 0.567 |
|  | R LTL-R AMY | 0.003 | 1.216 | 0.270 | -0.002 | 0.436 | 0.509 |
|  | R vlPFC-L HC | -0.005 | 5.536 | **0.019** | 0.001 | 0.257 | 0.612 |
|  | L LTL-L HC | 0.000 | 0.026 | 0.871 | -0.005 | 3.903 | 0.048 |
|  | L AMY-L HC | 0.003 | 1.206 | 0.272 | -0.008 | 6.517 | **0.011** |

**Table S8. Summary of reduced GLMEs of fronto-temporo-limbic coherence.** P-values were FDR-corrected for 43 reduced models of coherence with a critical value of p=0.005. Bolded p-values denote effects that survived FDR-correction.

**Table S9. Summary of Conflict x Group GLMEs of fronto-temporo-limbic power of participants included in DCM analyses.** P-values were FDR-corrected for 80 models of band power with a critical value of p=0.009. Bolded p-values denote effects that survived FDR-correction.

**Table S10. Summary of reduced GLMEs of fronto-temporo-limbic power of participants included in DCM analyses.** P-values were FDR-corrected for 64 reduced models of band power with a critical value of p=0.01. Bolded p-values denote effects that survived FDR-correction.

**Supplemental References**

1.	Oostenveld R, Fries P, Maris E, Schoffelen JM. FieldTrip: Open source software for advanced analysis of MEG, EEG, and invasive electrophysiological data. *Comput Intell Neurosci*. 2011;2011:156869. doi:10.1155/2011/156869

2.	Rosenberg JR, Amjad AM, Breeze P, Brillinger DR, Halliday DM. The Fourier approach to the identification of functional coupling between neuronal spike trains. *Prog Biophys Mol Biol*. 1989;53(1):1-31. doi:10.1016/0079-6107(89)90004-7

3.	*allfitdist*. MATLAB Central File Exchange; 2012.
